# “What” and “when” predictions modulate auditory processing in a contextually specific manner

**DOI:** 10.1101/2022.06.20.496917

**Authors:** Cappotto Drew, Luo Dan, Lai Hiu Wai, Peng Fei, Melloni Lucia, Schnupp Jan Wilbert Hendrik, Auksztulewicz Ryszard

## Abstract

Extracting regularities from ongoing stimulus streams to form predictions is crucial for adaptive behavior. Such regularities exist in terms of the content of the stimuli (i.e., “what” it is) and their timing (i.e., “when” it will occur), both of which are known to interactively modulate sensory processing. In real-world stimulus streams, regularities also occur contextually - e.g. predictions of individual notes vs. melodic contour in music. However, it is unknown whether the brain integrates predictions in a contextually congruent manner (e.g., if slower “when” predictions selectively interact with complex “what” predictions), and whether integrating predictions of simple vs. complex features rely on dissociable neural correlates. To address these questions, our study employed “what” and “when” violations at different levels - single tones (elements) vs. tone pairs (chunks) - within the same stimulus stream, while neural activity was recorded using electroencephalogram (EEG) in participants (N=20) performing a repetition detection task. Our results reveal that “what” and “when” predictions interactively modulated stimulus-evoked response amplitude in a contextually congruent manner, but that these modulations were shared between contexts in terms of the spatiotemporal distribution of EEG signals. Effective connectivity analysis using dynamic causal modeling showed that the integration of “what” and “when” prediction selectively increased connectivity at relatively late cortical processing stages, between the superior temporal gyrus and the fronto-parietal network. Taken together, these results suggest that the brain integrates different predictions with a high degree of contextual specificity, but in a shared and distributed cortical network.

**Significance statement:** Predictions of stimulus features, present in different statistically-regular contexts in the environment, are crucial to forming adaptive behavior. However, it is unknown if the brain integrates predictions selectively according to such contextual differences. By recording human electroencephalography during experimental manipulations of time-based and content-based predictions, we found that those predictions interactively modulated neural activity in a contextually congruent manner, such that local (vs. global) time-based predictions modulated content-based predictions of sequence elements (vs. chunks). These modulations were shared between contextual levels in terms of the spatiotemporal distribution of neural activity. This suggests that the brain integrates different predictions with a high degree of contextual specificity, but in a shared and distributed cortical network.

## Introduction

The ability to predict future events based on sensory information is an integral aspect of adaptive sensory processing. Real-world events are complex, consist of statistical regularities, and contain multiple features over which predictions can be formed (Dehaene et al., 2015). In the auditory domain, “what” and “when” predictions are present in virtually every stimulus stream, and their manipulation has been the foundation for numerous studies of predictive coding. “What” predictions are typically manipulated by introducing unexpected sensory deviants (oddballs), and comparing the neural responses to the unexpected vs. expected stimuli. In such oddball paradigms, the resulting classical mismatch response (MMR) is commonly interpreted as an error correction signal (Garrido et al., 2009). As opposed to “what” predictions, which often rely on MMR-based explanations, “when” predictions are typically explained by neural entrainment - phase alignment of neural activity to an external temporal structure (Auksztulewicz et al., 2019; Haegens and Zion Golumbic, 2018; Schroeder and Lakatos, 2009) (but see: (Doelling and Assaneo 2021)). An influential study (Ding et al., 2016) has suggested that cortical activity can selectively entrain to contextual structures in linguistic sequences pursuant to levels of chunking. More recently, this finding has been extrapolated to artificial streams of auditory and visual stimuli (Henin et al., 2021).

Several studies have investigated predictions through independent manipulation of timing and content predictability, suggesting interactive and partly dissociable neural correlates and putative underlying mechanisms (Kotz & Schwartze, 2010; Arnal & Giraud, 2012; Buzsaki & Friston, 2016; Auksztulewicz et al., 2018). MMR amplitudes are typically modulated by “when” predictions, such that deviant-evoked activity is higher when deviants are presented in temporally predictable (e.g., rhythmic/isochronous) sequences (Jalewa et al., 2021; Lumaca et al., 2019; Takegata and Morotomi, 1999; Todd et al., 2018; Yabe et al., 1997). In the auditory domain, such interactions have been suggested to rely on partially dissociable networks (Hsu, et al., 2013), while also jointly modulating stimulus-evoked activity in the superior temporal gyrus (Auksztulewicz et al., 2018). More generally, it has been proposed that interactions between “what” and “when” predictions are inherent to the processing of musical sequences (Musacchia et al., 2014). In this context, it has been suggested that neural entrainment along the non-lemniscal (secondary) auditory pathway (sensitive to the rhythmic sequence structure) can modulate activity in the lemniscal (primary) pathway (encoding stimulus contents), including MMR processing.

However, it is unknown if interactions between “what” predictions (in the lemniscal pathway) and “when” predictions (in the non-lemniscal pathway) are specific to differing contexts present in complex naturalistic stimuli such as speech or music (Hasson et al., 2015). In the case of naturalistic music stimuli, lower-level predictions can be formed about single notes within a sequence, while higher-level predictions can relate to the resulting melody contour, each occurring at their respective time scales. In principle, neural entrainment to a particular time scale might boost the processing of any stimuli presented in the expected time window (Auksztulewicz et al., 2019). However, if entrainment to slower (i.e., more global) temporal scales is functionally related to chunking (Ding et al., 2016; Henin et al., 2021), it may show a specific modulation of the processing of stimulus chunks, rather than single elements. Thus, based on current hypotheses of neural entrainment and predictive processing, it is unclear if “when” predictions modulate the processing of stimulus contents (and the respective “what” predictions) in a contextually specific way - e.g. if temporal predictions amplify the processing of any stimuli presented at a preferred time window, or only those stimuli whose contents can be predicted at the corresponding time scale.

Here, we present streams of tones and independently manipulate content-based and time-based characteristics of the stream at two levels, while recording EEG in healthy volunteers. Temporal predictability was manipulated at slower (∼2 Hz) and faster (∼4 Hz) time scales, while acoustic deviants were introduced at lower (e.g. single tones) and higher (e.g.chunked tone pairs) levels, to evaluate the independent or interactive effect of “what” and “when” predictive processing across contexts.

## Methods

EEG was recorded during an auditory repetition detection task in order to gauge (1) the effects of “when” predictions at higher and lower temporal scales on tone-evoked responses and on neural entrainment, as well as (2) the modulatory effect of “when” predictions on the neural signatures of higher and lower-level “what” predictions (MMRs). The use of musical sequences (ascending or descending musical scales) was chosen to reduce the influence of speech-specific processing on neural activity (e.g., modulation by language comprehension, speech-specific semantic and syntactic processing, etc.) and provide a better comparison to similar work in animal models (Jalewa et al., 2021). In the analysis, we focused on interactions between “what” and “when” predictions, specifically testing whether MMRs are modulated by temporal predictability in a contextually specific way (such that slower “when” predictions selectively modulate MMRs to violations of higher-level “what” predictions). To explain the effects observed at the scalp level, we used source reconstruction and biophysically realistic computational modeling (dynamic causal modeling), which allowed us to infer the putative mechanisms of interactions between “what” and “when” predictions.

### Participant sample

Participants (N=20, median age 21, range 19-25), 10 females, 10 males; 19 right-handed, 1 left-handed) volunteered to take part in the study upon written consent. The work was conducted in accordance with protocols approved by the Human Subjects Ethics Sub-Committee of the City University of Hong Kong. All participants self-reported normal hearing and no current or past neurological or psychiatric disorders.

### Stimulus design and behavioral paradigm

An experimental paradigm was designed in which auditory sequences were manipulated with respect to “what” and “when” predictions at two contextual levels (“what” predictions of single tones vs. chunked tone pairs; “when” predictions at ∼4 Hz vs. ∼2 Hz), allowing for an analysis of their interactions at each level. To ensure that participants paid attention to stimulus sequences, the sequences contained very occasional repetitions, and participants were instructed to listen out for such repetitions (see below). The experimental manipulations of “what” and “when” predictions, however, were irrelevant to the task, such that neural responses to unexpected stimuli are not confounded by neural activity related to target detection.

Auditory sequences were generated using Psychtoolbox for MATLAB (version 2021a) and delivered to participants fitted with Brainwavz B100 earphones via a TDT RZ6 multiprocessor at a playback sampling rate of 24414 Hz. Participants were seated in a sound-attenuated EEG booth. Visual stimuli (fixation cross) and instructions were presented on a 24-inch computer monitor and delivered using the Psychophysics Toolbox for MATLAB. Participants were asked to minimize movements and eye blinks and instructed to perform a tone repetition detection task, by pressing a keyboard button using their right index finger as soon as possible upon hearing an immediate tone repetition.

Stimuli were presented in sequences of 7 ascending or descending scales. Each scale was composed of 8 tones equally spaced on a logarithmic scale to form one octave. Thus, across 7 scales a total 56 tones were presented per sequence (Fig. 1A, 1B). A trial was defined as the presentation of a sequence of 7 scales. Within a trial, all scales were either ascending or descending. The ascending and descending trials were presented in a random order. Each participant heard a total of 240 sequences (trials). The initial tone of each scale was randomly drawn from a frequency range 300-600 Hz. Each tone was generated by resynthesizing a virtual harp note F4 (played on virtualpiano.net), to match a fixed 166 ms duration and the fundamental frequency used at a given position in the scale. The tone manipulations were implemented in an open-source vocoder, STRAIGHT (Kawahara, 2006) for Matlab 2018b (MathWorks; RRID: SCR_001622). Tones were perceptually grouped into pairs by manipulating the intensity ratio of odd/even tones, with the even (2nd, 4th, 6th and 8th) tones within a scale presented 10 dB quieter relative to the odd-position tones (Kotz et al., 2018).

**Figure 1.**
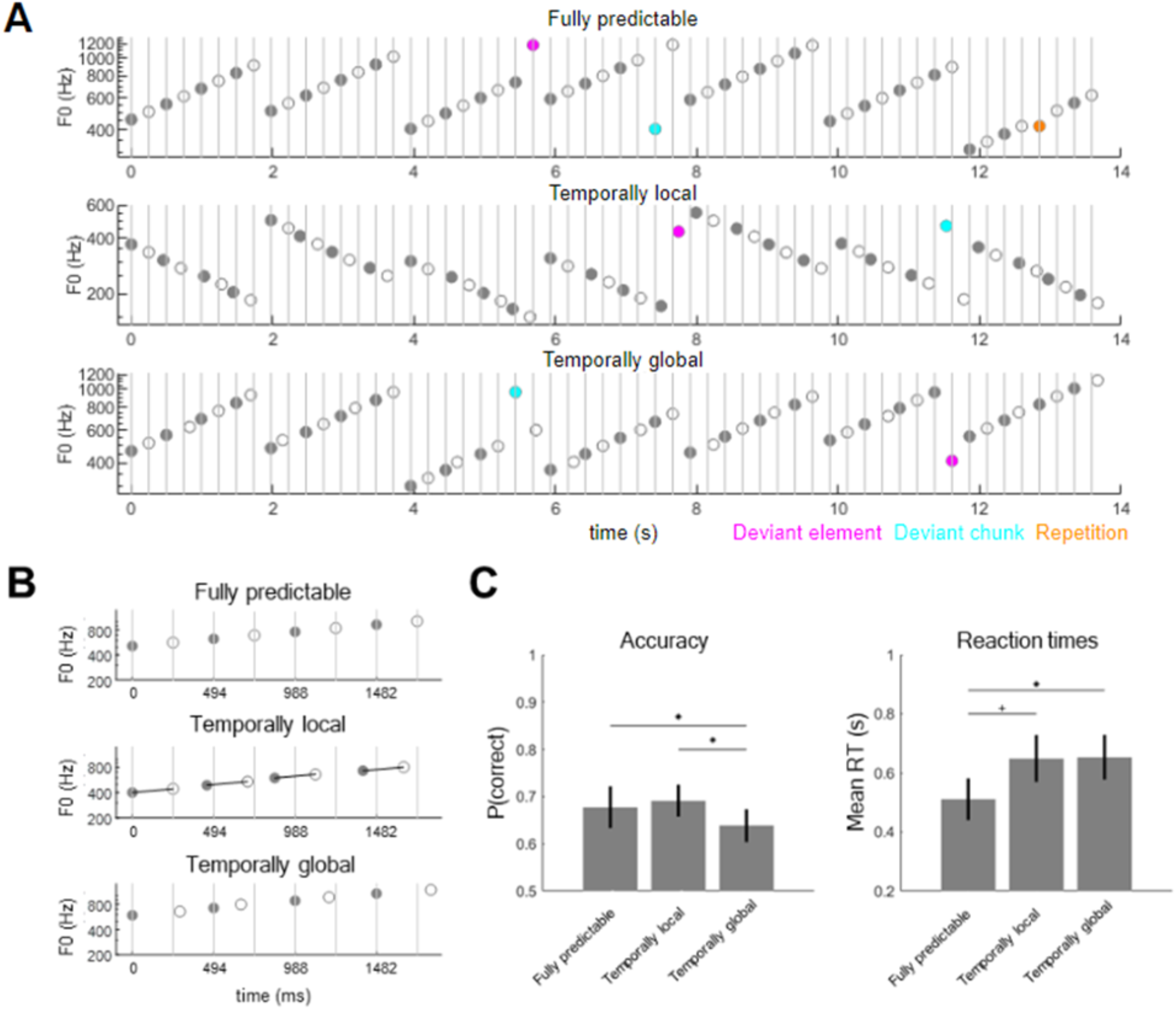
Experimental paradigm and behavioral results. **(A)** Participants listened to sequences of ascending (as represented on the figure) or descending scales of acoustic tones. Sequences were composed of tone pairs, where odd tones (gray circles) were louder than even tones (white circles). Participants performed a tone repetition detection task (orange circles: behavioral targets; presented in a subset of trials). Additionally, sequences could include deviant tones (magenta circles), in which one of the pair-final tones had an outlier fundamental frequency (F0), and deviant chunks (cyan circles), in which one of the pair-initial tones had an outlier F0. **(B)** Sequences were blocked into three temporal conditions: a fully-predictable condition (upper panel), in which ISI between tones was fixed at 0.247 s; a temporally-local condition (middle panel), in which the ISI between odd and even tones within pairs was fixed at 0.247 s but the ISI between odd tones (pair-initial tones) was jittered; and a temporally-global condition (lower panel), in which ISI between odd tones (pair-initial tones) was fixed at 0.494 s but the ISI between odd and even tones within pairs was jittered. **(C)** Behavioral results. Left panel: accuracy, right panel: reaction times. Error bars denote SEM across participants. Asterisks denote p < 0.05, plus symbol denotes a trend towards significance.

Manipulation of temporal predictability formed three conditions: in the fully-predictable (isochronous) condition, tones were presented with a fixed ISI (inter-stimulus interval) of 247 ms, resulting in all tones having predictable timing at both the slow time scale (chunks) and the fast time scale (elements). In the temporally-global (predictable slow, unpredictable fast) condition, the slow time scale was predictable (corresponding to a fixed pair onset asynchrony, i.e., a fixed 494 ms interval between the onsets of the odd, pair-initial tones) but the fast time scale was unpredictable (corresponding to a random onset of the even, pair-final tones, relative to the pair-initial tones). In this condition, the exact ISI of the pair-final tones was set by randomly drawing one value from the following 4 ISIs, relative to the standard 247 ms ISI: 33.3% shorter; 16.6% shorter; 16.6% longer; 33.3% longer. Finally, in the temporally-local (predictable fast, unpredictable slow) condition, the onset of stimuli at the fast time scale was predictable (corresponding to a fixed 247 ms ISI of the pair-final tones, relative to the pair-initial tones) but the slow time scale was unpredictable (corresponding to a random onset of the odd, pair-initial tones, relative to the expected 494 ms interval). In this condition, the exact ISI of the pair-initial tones was set by randomly drawing one value from the same 4 ISIs as above, and shifting the onset of the pair-initial tone by this value, relative to the expected 494 ms interval relative to the previous pair onset. A fixed inter-trial interval of 1 second was employed between the offset of the last tone of a 56-tone sequence and the onset of the first tone in the next sequence. The three timing conditions were administered in 12 blocks of 20 trials (4 blocks per condition). Blocks were pseudo-random in order, allowing no immediate repetitions of the same, timing, condition.

Content predictability was manipulated by altering the fundamental frequency of a subset of tones within the scales, such that trials could contain an element deviant (i.e., a single deviant tone) or a “chunk” deviant (i.e., a deviant tone pair). The element deviants were introduced by replacing the final tone of a scale with an outlier frequency (i.e., a tone whose fundamental frequency was 20% lower/higher than the range of the entire scale). The chunk deviants were introduced by replacing the penultimate tone of a scale (i.e., the initial tone of the final pair, rendering the entire pair unpredictable) in the same manner.

To facilitate the extraction of statistical regularities in the sequences, in each trial, the first two scales were left unaltered. Two deviant tones were randomly placed within the subsequent 5 scales. Additionally, in 50% of the trials, a scale containing an immediate tone repetition was included in the last 5 scales. In subsequent EEG analysis, neural responses evoked by element and chunk deviants were compared with neural responses evoked by the respective standard tones, designated as the final (standard element) and penultimate (standard chunk) tones in two unaltered scales out of the final 5.

In total, 64.3% of the scales were left unaltered, 14.3% contained an element deviant, 14.3% contained a chunk deviant, and 7.1% contained a tone repetition. The global deviant probability equaled 3.57% of all tones, amounting to 80 deviant tones per deviant type (element, chunk) per temporal condition (fully predictable, temporally local, temporally global). To ensure that the EEG analysis is not confounded by differences in baseline duration between temporal conditions (e.g., element deviants preceded by shorter/longer ISIs in the temporally global condition than in the other two conditions), the ISIs preceding all deviant tones and designated standard tones were replaced by a fixed 247 ms ISI. Therefore, the temporal predictability manipulation was limited to tones surrounding the analyzed tones and did not affect the exact timing of either deviants or standards.

Prior to experimental blocks, participants were exposed to a training session consisting of fully predictable sequences containing a tone repetition, to familiarize themselves with the task and stimuli. Participants performed training trials until they could detect tone repetition in 3 consecutive trials with reaction times shorter than 2 seconds. Then, during the actual experiment, participants received feedback on their mean accuracy and reaction time after each block of 20 trials. The data segments (scales) containing tone repetition were subsequently discarded from EEG analysis.

### Behavioral analysis

Analysis was performed on the accuracy and reaction time data corresponding to participant responses during the repetition detection task. Reaction times longer than 2 seconds were excluded from analysis. Mean reaction times (from correct trials only) were log-transformed to approximate a normal distribution. Accuracy and mean reaction times were entered into separate repeated-measures ANOVAs with a within-subjects factor Time (fully predictable, temporally local, temporally global). Post-hoc comparisons were implemented using paired t-tests in MATLAB.

### Neural data acquisition and pre-processing

EEG signals were collected using a 64-channel ANT Neuro EEGo Sports amplifier at a sampling rate of 1024 Hz with no online filters. The recorded data were pre-processed using the SPM12 Toolbox (version 7219; Wellcome Trust Centre for Neuroimaging, University College London; RRID: SCR_007037) for MATLAB (version R2018b). Continuous data were high-pass filtered at 0.1 Hz and notch filtered between 48 Hz and 52 Hz before being down-sampled to 300 Hz and subsequently low-pass filtered at 90 Hz. All filters were 5th order zero-phase Butterworth. Eyeblink artifacts were detected using channel Fpz and removed by subtracting the two top spatiotemporal principal components of eyeblink-evoked responses from all EEG channels (Ille et al., 2002). Cleaned signals were re-referenced to the average of all channels, as is recommended for source reconstruction and dynamic causal modeling (Litvak et al. 2011). The pre-processed data were analyzed separately in the frequency domain (phase coherence analysis) and in the time domain (event-related potentials; ERPs).

### Phase coherence analysis

To test whether tone sequences are associated with dissociable spectral peaks in the neural responses at the element rate (4.048 Hz) and at the chunk rate (2.024 Hz), we analyzed the data in the frequency domain. Continuous data were segmented into epochs ranging from the onset to the offset of each trial (tone sequence). For each participant, channel, and sequence, we calculated the Fourier spectrum of EEG signals measured during that sequence. Based on previous literature, we then calculated the inter-trial phase coherence (ITPC), separately for each temporal condition (fully-predictable, temporally-local, temporally-global) according to the following equation (Ding and Simon, 2013) in order to infer phase consistency in each condition:

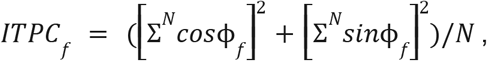

where *φ*_*f*_ corresponds to the Fourier phase at a given frequency *f*, and *N* corresponds to the number of sequences (80 per condition). The same method was used to estimate the stimulus frequency spectrum by calculating the ITPC based on the raw stimulus waveform.

In the initial analysis, ITPC estimates were averaged across EEG channels. To test for the presence of statistically significant spectral peaks, ITPC values at the single-tone rate (4.048 Hz) and tone-pair rate (2.024 Hz) were compared against the mean of ITPC values at their respective neighboring frequencies (single-tone rate: 3.974 and 4.124 Hz; tone-pair rate: 1.949 and 2.099 Hz) using paired t-tests.

Furthermore, to test whether element-rate and chunk-rate spectral peaks observed at single EEG channels show modulations due to temporal predictability, spatial topography maps of single-channel ITPC estimates were converted to 2D images, smoothed with a 5 × 5 mm full-width-at-half-maximum (FWHM) Gaussian kernel, and entered into repeated-measures ANOVAs (separately for element-rate and chunk-rate estimates) with a within-subjects factor Time (fully predictable, temporally local, temporally global), implemented in SPM12 as a general linear model (GLM). To account for multiple comparisons and for ITC correlations across neighboring channels, statistical parametric maps were thresholded at p < 0.001 and corrected for multiple comparisons over space at a cluster-level p_FWE_ < 0.05 under random field theory assumptions (Kilner et al., 2005).

Finally, to test whether spectral signatures of temporal predictability are modulated by experience with stimuli, we split the data into two halves (two consecutive bins of 40 trials), separately for each condition. Element-rate and chunk-rate ITPC estimates were averaged across EEG channels and compared separately for each of the two halves using repeated-measures ANOVAs with a within-subjects factor Time (fully predictable, temporally local, temporally global).

### Event-related potentials

For the time-domain analysis, data were segmented into epochs ranging from -50 ms before to 247 ms after deviant/standard tone onset, baseline-corrected from -25 ms to 25 ms to prevent epoch contamination due to the temporally structured presentation (Fitzgerald et al., 2021), and denoised using the “Dynamic Separation of Sources” (DSS) algorithm (de Cheveigné and Simon, 2008). Condition-specific ERPs (corresponding to element/chunk deviants and the respective standards, presented in each of the three temporal conditions) were calculated using robust averaging across trials, as implemented in the SPM12 toolbox, and low-pass filtered at 48 Hz (5th order zero-phase Butterworth). The resulting ERPs were analyzed univariately to gauge the effects of “what” and “when” predictions on evoked responses. ERP data were converted to 3D images (2D: spatial topography; 1D: time), and the resulting images were spatially smoothed using a 5 × 5 mm FWHM Gaussian kernel. The smoothed images were entered into a general linear model (GLM) implementing a 3 × 3 repeated-measures ANOVA with a within-subject factors Contents (standard, deviant element, deviant chunk) and Time (fully predictable, temporally local, temporally global). Beyond testing for the two main effects and a general 3 × 3 interaction, we also designed a planned contrast quantifying the congruence effect (i.e., whether “when” predictions specifically modulate the amplitude of mismatch signals evoked by deviants presented at a time scale congruent with “when” predictions, i.e., deviant elements in the temporally-local condition and deviant chunks in the temporally-global conditions). To this end, we tested for a 2 × 2 interaction between Contents (deviant element, deviant chunk) and Time (temporally local, temporally global). To account for multiple comparisons as well as for ERP amplitude correlations across neighboring channels and time points, statistical parametric maps were thresholded at p < 0.001 and corrected for multiple comparisons over space and time at a cluster-level p_FWE_ < 0.05 under random field theory assumptions (Kilner et al., 2005).

### Brain-behavior correlations

To test whether the neural effects of “what” and/or “when” predictive processing correlate with each other, as well as with behavioral benefits of “when” predictions in the repetition detection task, we performed a correlation analysis across participants. Thus, for each participant, we calculated a single behavioral index (the difference between accuracy scores obtained in the temporally local vs. temporally global condition) and three statistically significant neural indices. The first neural index - the “congruence effect” - quantified the difference between deviant-evoked ERP amplitudes measured in the temporally congruent (deviant elements presented in the temporally local condition; deviant chunks presented in the temporally global condition) and incongruent (deviant chunks presented in the temporally local condition; deviant elements presented in the temporally global condition) conditions, averaged across electrodes in the significant cluster where we observed a significant congruence effect (i.e., a 2 × 2 interaction between “what” and “when” predictions; see Results and Fig. 3C). The second neural index - the “ITPC effect” - quantified the difference between the chunk-rate ITPC values obtained for temporally local vs. temporally global conditions in the second half of the experiment (see Results Fig. 2D). The third neural index - the “mismatch effect” - quantified the difference between the absolute deviant-evoked and standard-evoked ERP amplitudes (averaged across significant channels and temporal conditions; Fig. 3AB), since we hypothesized that performance in the repetition detection task might be related to overall deviance detection, we also included an index of “what” predictions. We then fitted a linear regression model with three predictors (i.e., the three neural indices) regressed against the behavioral accuracy index, and identified outlier participants using a threshold of Cook’s distance exceeding 5 times the mean. Correlations between all measures were quantified using Pearson’s *r* and corrected for multiple comparisons using Bonferroni correction, implementing a conservative correction given no a priori assumptions about the correlation coefficients.

**Figure 2.**
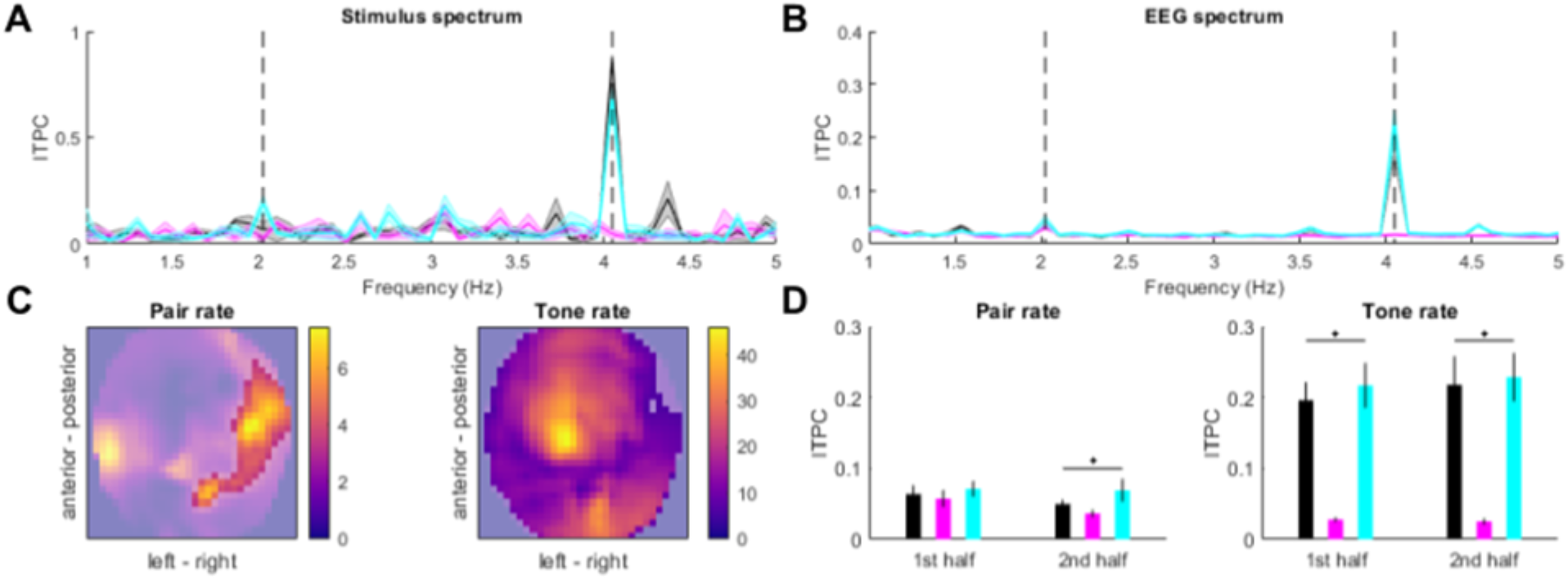
Spectral signatures of temporal predictability. (A) Inter-trial phase coherence (ITPC) in the stimulus spectrum. Black: fully-predictable, cyan: temporally-local, magenta: temporally-global. Chunk-rate (2.024 Hz) and element-rate (4.048 Hz) peaks are indicated by dashed vertical lines. (B) ITPC based on EEG activity (averaged across channels). Legend as above. Shaded areas indicate SEM across participants. (C) EEG topography maps of main effects of Condition (fully predictable vs. temporally local vs. temporally global) on the chunk-rate peak ITPC (left panel) and tone-rate peak ITPC (right panel). Statistical F values are represented on the color scale. Unmasked area corresponds to significant clusters (p_FWE_ < 0.05). (D) Chunk-rate (left panel) and element-rate (right panel) peak ITPC values plotted separately for the 1st half and 2nd half of the trials. Error bars denote SEM across participants.

**Figure 3.**
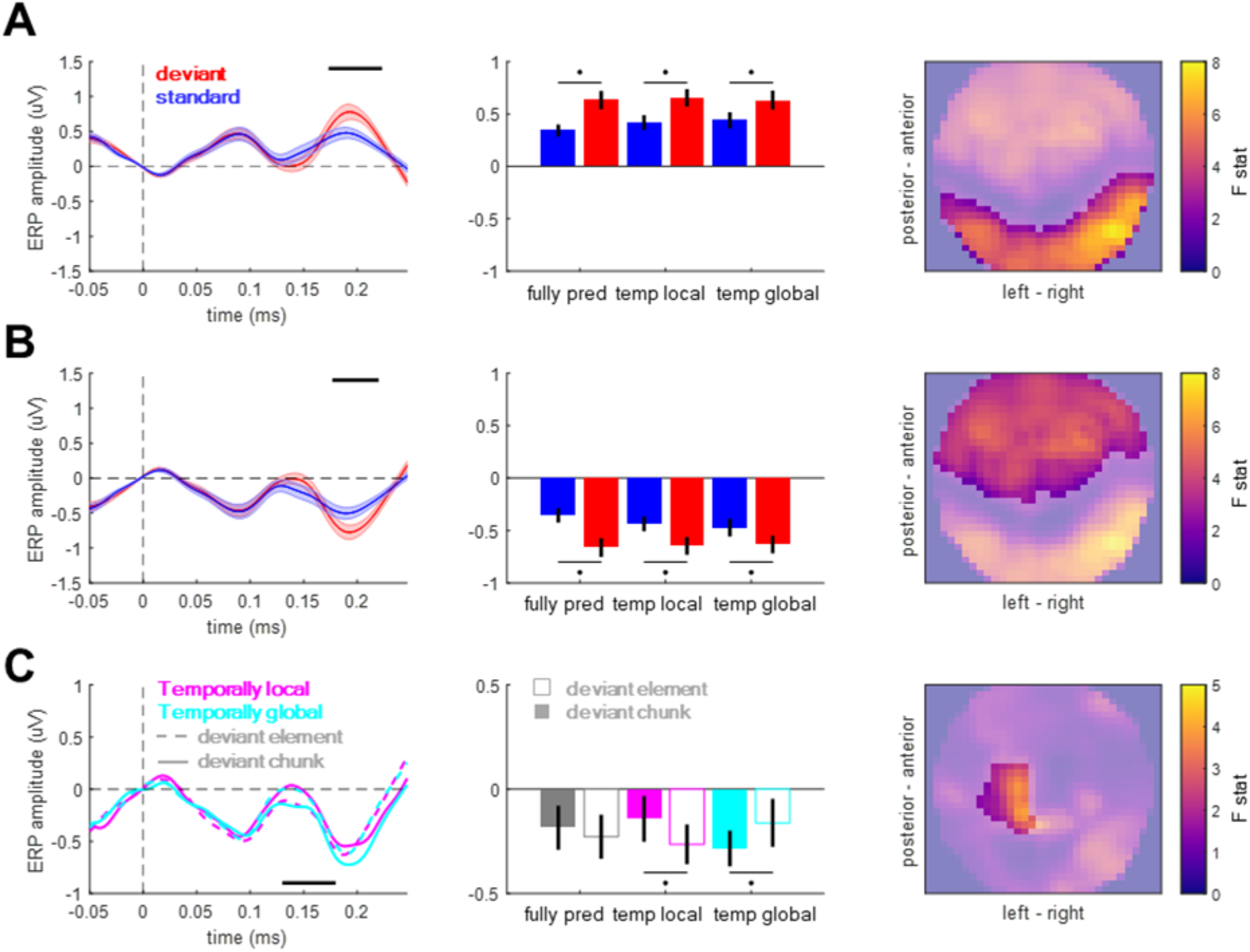
Event-related potentials. **(AB)** Main effect of content-based predictions (deviant vs. standard) in anterior (A) and posterior (B) clusters. Left panels: time courses of ERPs averaged over the spatial topography clusters shown in the right panels. Shaded area denotes SEM across participants. Black horizontal bar denotes p_FWE_ < 0.05. Middle panels: mean voltage values for standards (blue) and deviants (red). Right panels: spatial distribution of the main effect. Color bar: F value. **(C)** Contextual interaction between content-based predictions (deviant element vs. deviant chunk) and temporal predictions (temporally global vs. temporally local). Left panels: time courses of ERPs averaged over the spatial topography clusters shown in the right panels. Black horizontal bar denotes p_FWE_ < 0.05. Middle panels: mean voltage values for the six deviant conditions. Right panels: spatial distribution of the interaction effect. Color bar: F value.

### Source reconstruction

Source reconstruction was performed under group constraints (Litvak and Friston, 2008) which allows for an estimation of source activity at a single-participant level under the assumption that activity is reconstructed in the same subset of sources for each participant. Sources were estimated using empirical Bayesian beamformer (Belardinelli et al., 2012; Little et al., 2018; Wipf and Nagarajan, 2009) based on the entire post-stimulus time window (0-247 ms). Since in the ERP analysis (see Results) we identified two principal findings - namely a difference between ERPs evoked by deviants and standards, and an interaction between deviant type and temporal condition - we focused on comparing source estimates corresponding to these effects. In the analysis of the difference between deviants and standards, source estimates were extracted for the 173-223 ms time window, converted into 3D images consisting of 3 spatial dimensions and smoothed with a 10 × 10 × 10 mm FWHM Gaussian kernel. Smoothed images were then entered into a GLM implementing a 3 × 3 repeated-measures ANOVA with within-subjects factors of Content (standard, deviant element, deviant chunk) and Time (fully predictable, temporally local, temporally global). In the analysis of the interaction between deviant type and temporal condition, source estimates were extracted for the 130-180 ms and processed as above. Smoothed images were then entered into a GLM implementing a 2 × 2 repeated-measures ANOVA with within-subjects factors of Content (deviant element, deviant chunk) and Time (temporally local, temporally global). To account for multiple comparisons as well as for source estimate correlations across neighboring voxels, statistical parametric maps were thresholded and corrected for multiple comparisons over space at a cluster-level p_FWE_ < 0.05 under random field theory assumptions (Kilner et al., 2005). Source labels were assigned using the Neuromorphometrics probabilistic atlas, as implemented in SPM12.

### Dynamic causal modeling

Dynamic causal modeling (DCM) was used to estimate source-level connectivity parameters associated with general mismatch processing (deviant vs. standard) and with the contextual interaction between “what” and “when” predictions (element deviant presented in the temporally-local condition, and chunk deviant presented in the temporally-global condition, vs. element deviant presented in the temporally global condition, and chunk deviant presented in the temporally-local condition). DCM is a type of an effective connectivity analysis based on a generative model, which maps the data measured at the sensor level (here: EEG channels) to source-level activity. The generative model comprises a number of sources which represent distinct cortical regions, forming a sparse interconnected network. Activity in each source is explained by a set of neural populations, based on a canonical microcircuit (Bastos et al., 2012), and modeled using coupled differential equations that describe the changes in postsynaptic voltage and current in each population. Here, we used a microcircuit consisting of four populations (superficial and deep pyramidal cells, spiny stellate cells, and inhibitory interneurons), each having a distinct connectivity profile of ascending and descending extrinsic connectivity (linking different sources) and intrinsic connectivity (linking different populations within each source). The exact form of the canonical microcircuit and the connectivity profile was identical as in previous literature on the topic (Auksztulewicz et al., 2018; Auksztulewicz and Friston, 2015; Fitzgerald et al., 2021; Rosch et al., 2019; Todorovic and Auksztulewicz, 2021). Importantly for our study, a subset of intrinsic connections corresponds to self-connectivity parameters, describing the neural gain of each region. Both extrinsic connectivity and gain parameters were allowed to undergo condition-specific changes, modeling differences between experimental conditions (deviants vs. standards, and the hierarchical interaction between “what” and “when” predictions).

Here, we used DCM to reproduce the single-participant, condition-specific ERPs in the 0-247 ms range. Based on the source reconstruction (see Results) and previous literature (Garrido et al., 2009), we included six sources in the cortical network: bilateral primary auditory cortex (A1; Montreal Neurological Institute coordinates: left, [-42 -22 7]; right, [46 -14 8]), bilateral superior temporal gyrus (STG; left, [-60 -20 -8]; right, [59 -25 8]), right inferior frontal gyrus (IFG; [40 26 -6]), and left superior parietal lobule (SPL; [-26 -40 46]). To quantify model fits, we used the free-energy approximation to model evidence, penalized by model complexity. The analysis was conducted in a hierarchical manner-first, model parameters (including extrinsic and intrinsic connections, as well as their condition-specific changes) were optimized at the single participants’ level, and then the significant parameters were inferred at the group level.

At the first level, models were fitted to single participants’ ERP data over two factors: “what” predictions (all deviants vs. standards) and the contextual interaction between “what” and “when” predictions (element deviant presented in the temporally-local condition, and chunk deviant presented in the temporally-global condition, vs. element deviant presented in the temporally-global condition, and chunk deviant presented in the temporally-local condition). At this level, all extrinsic and intrinsic connections were allowed to be modulated by both factors, corresponding to a “full” model.

Since model inversion in DCM is susceptible to local maxima due to the inherently nonlinear nature of the models, the analysis at the second (group) level implemented parametric empirical Bayes (Friston et al., 2015). Therefore, group-level effects were inferred by (re)fitting the same “full” models to single participants’ data, under the assumption that model parameters should be normally distributed in the participant sample, and updating the posterior distribution of the parameter estimates. We used Bayesian model reduction (Friston and Penny, 2011) to compare the “full” models against a range of “reduced” models, in which some parameters were not permitted to be modulated by the experimental factors. Specifically, we designed a space of alternative models, such that each model allowed for a different subset of connections to contribute to the observed ERPs. The model space examined each combination of modulations of (1) ascending connections (e.g., from A1 to STG), (2) descending connections (e.g., from STG to A1), (3) lateral connections (e.g., from left to right STG), and (4) intrinsic connections (i.e., gain parameters). This resulted in 256 models (16 models for each of the two factors). The free-energy approximation to log-model evidence was used to score each model. Since no single winning model was selected (see Results), Bayesian model averaging was used to obtain weighted averages of posterior parameter estimates, weighted by the log-evidence of each model. This procedure yielded Bayesian confidence intervals for each parameter, quantifying the uncertainty of parameter estimates. Parameters with 99.9% confidence intervals falling either side of zero (corresponding to p < 0.001) were selected as statistically significant.

## Results

### Behavioral results

Performance across all trials revealed significant differences in accuracy across conditions (main effect of Time: F_2,38_ = 7.3530, p = 0.002), corresponding to significantly lower accuracy in the temporally-global condition (mean ± SEM: 63.88% ± 3.65%) than in the fully-predictable (mean ± SEM: 67.75% ± 4.64%; t19 = -2.5272, p = 0.0205) and temporally-local conditions (mean ± SEM: 69.12% ± 3.55%; t19 = -5.984, p < 0.001) (Fig. 1C). Reaction times also significantly differed across conditions (F_2,38_ = 3.5543, p = 0.0385), with post-hoc analysis revealing that reaction times were significantly faster in the fully-predictable condition (mean ± SEM: 511 ± 74 ms) than in the temporally-global condition (mean ± SEM: 653 ± 79 ms; t19 = 2.4089, p = 0.0263). The difference between the fully-predictable condition and the temporally-local condition (mean ± SEM: 649 ± 83 ms) trended towards significance (t19 = 2.0132, p = 0.0585). No significant difference was observed between the temporally-global condition and the temporally-local condition (p = 0.9013).

### Phase coherence analysis

In the EEG spectrum of inter-trial phase coherence (ITPC; averaged across conditions and channels), both element-rate peak (4.048 Hz) and chunk-rate peak (2.024 Hz) were observed, relative to neighboring frequency points (element-rate: t_19_ = 6.8489, p < 0.001; chunk-rate: t_19_ = 3.6274, p = 0.0018). The ITPC peak estimates differed between experimental conditions, reflecting differences in the stimulus spectrum. Specifically, the chunk-rate ITPC estimates were higher in the fully-predictable and temporally-global conditions than in the temporally-local conditions, and this effect was observed at most of the EEG channels (F_max_ = 46.30, Z_max_ = 6.43, p_FWE_ < 0.001; pairwise comparisons: fully-predictable vs. temporally-local, T_max_ = 8.02, Z_max_ = 6.10, p_FWE_ < 0.001; temporally-global vs. temporally-local, T_max_ = 9.62, Z_max_ = 6.81, p_FWE_ < 0.001; fully-predictable vs. temporally-global, all p_FWE_ > 0.05). On the other hand, the chunk-rate ITPC estimates were higher in the temporally-global condition than in the other two conditions, and this effect was observed over right lateral channels (F_max_ = 7.45, Z_max_ = 2.90, p_FWE_ = 0.031; pairwise comparisons: temporally-global vs. fully-predictable, T_max_ = 3.81, Z_max_ = 3.48, p_FWE_ = 0.004; temporally-global vs. temporally-local, T_max_ = 3.83, Z_max_ = 3.50, p_FWE_ = 0.001; fully-predictable vs. temporally-local, all p_FWE_ > 0.05). Interestingly, the chunk-rate differences between conditions built up during the experiment: they were absent during the first half of the experiment (F_2,59_ = 1.0433, p = 0.3622), and were only observed during the second half of the experiment (F_2,59_ = 3.8798, p = 0.0293). This was not the case for the element-rate differences between conditions, which were stable during the experiment (first half: F_2,59_ = 26.1701, p < 0.001; second half: F_2,59_ = 26.9480, p < 0.001).

The emergence of chunk-rate differences in ITPC over the course of the experiment was reflected in behavior. Specifically, RTs decreased for the second half of the experiment, relative to the first half, only for the temporally-global condition (Wilcoxon’s signed rank test: Z_19_ = -2.0926, p = 0.0364) but not for the fully-predictable condition (Z_19_ = -1.6902, p = 0.0910) or the temporally-local condition (Z_19_ = -0.8213, p = 0.4115). No differences in accuracy were observed for any of the three conditions across the first and second halves of the experiment (all p > 0.05).

### Event-related potentials

To test for effects of “what” and “when” predictions on ERP amplitudes, we analyzed the data in the time domain. ERP amplitudes differed significantly between deviant and standard tones, pooled over temporal conditions (Fig. 3A; posterior cluster: 173-223 ms, F_max_ = 53.94, Z_max_ = 6.68, p_FWE_ < 0.001; anterior cluster: 177-220 ms; F_max_ = 37.57; Z_max_ = 5.67; p_FWE_ < 0.001), corresponding to a typical anterior-posterior MMN topography after common-average referencing (Mahajan, Peter, and Sharma 2017). When analyzing specific deviant types (element and chunk deviants vs. their respective standards), significant differences between deviants and standards were observed in both cases (element deviants vs. standards: posterior cluster, 173-223 ms, F_max_ = 41.50, Z_max_ = 5.94, p_FWE_ < 0.001; anterior cluster, 177-227 ms; F_max_ = 35.56; Z_max_ = 5.52; p_FWE_ < 0.001; chunk deviants vs. standards: posterior cluster, 170-220 ms, F_max_ = 45.63, Z_max_ = 6.20, p_FWE_ < 0.001; anterior cluster, 177-213 ms; F_max_ = 30.17; Z_max_ = 5.11; p_FWE_ < 0.001). No significant differences were observed between the two deviant types, pooling over temporal conditions (p_FWE_ > 0.05). Thus, the main effect of “what” predictions differentiated between deviants and standards, but not between deviant types.

In the analysis of the main effect of “when” predictions (pooled over deviants and standards), no significant differences between the three temporal conditions were revealed (all p_FWE_ > 0.05). Similarly, in the analysis of the interaction effect of “what” and “when” predictions (pooled over deviant types), no significant effects were revealed. Specifically, neither deviants nor standards showed significant ERP amplitude differences when presented in different temporal contexts (all p_FWE_ > 0.05). Thus, the overall temporal structure of the sound sequences did not affect the element-evoked responses (averaged across deviants and standards) or the mismatch responses (differences between deviants and standards).

However, an analysis of the interaction between “what” and “when” predictions based on deviants presented in congruent temporal contexts (e.g. element deviants in the temporally-local condition) and those presented in non-temporally congruent contexts (e.g. element deviants in the temporally-global condition) revealed a significant interaction between deviant type and temporal condition (Fig. 3B; left central-posterior cluster: 130-180 ms, F_max_ = 20.63, Z_max_ = 4.24, p_FWE_ = 0.044). Post-hoc analysis revealed that MMR amplitudes in temporally-local were significantly larger for deviant elements (mean ± SEM: -0.1640 ± 0.0942 μV) than for deviant chunks (mean ± SEM: 0.0091 ± 0.1010 μV; t_19_ = 2.2843, p = 0.0340, two-tailed). In the temporally-global condition, MMR amplitude was observed to be nominally larger for deviant chunks (mean ± SEM: -0.1725 ± 0.0851 μV) than for deviant elements (mean ± SEM: -0.0155 ± 0.1233 μV), although the effect did not reach significance (t_19_ = 1.9024, p = 0.0724, two-tailed). No significant interaction effects were revealed when comparing deviant types between the fully-predictable condition and either the temporally-global or the temporally-local conditions. Thus, we observed a specific increase in deviant ERP amplitude when this deviant was presented in a temporally congruent context.

### Brain-behavior correlation analysis

Three neural predictors (the “congruence index”, quantifying the interactive effects of “what” and “when” predictions on ERPs; the “ITPC index”, quantifying the effect of “when” predictions on ITPC; and the “mismatch index”, quantifying the effect of “what” predictions on ERPs) were tested as potential correlates of the behavioral benefits in the repetition detection task accuracy. We identified two outlier participants based on a linear regression model. Having excluded these two participants, we did not find any significant correlations between the neural indices and the behavioral index (Pearson’s *r;* all p > 0.2), suggesting that behavior in the repetition detection task is not functionally related to ERP signatures of deviance detection. However, we did find a significant correlation between the congruence index and the ITPC index (r = 0.6439; p = 0.0039; corrected), such that the magnitude of the ERP difference between deviants presented in the temporally congruent vs. incongruent conditions positively correlated with the magnitude of the ITPC difference between temporally-local and temporally-global conditions.

### Source reconstruction

To identify the most plausible sources underlying the observed ERP differences between deviants and standards, as well as the contextual interaction between deviant types and temporal conditions, we carried out a source reconstruction analysis (Fig.4). Overall, source reconstruction explained 76.43 ± 3.08% (mean ± SEM across participants) of sensor-level variance.

**Figure 4.**
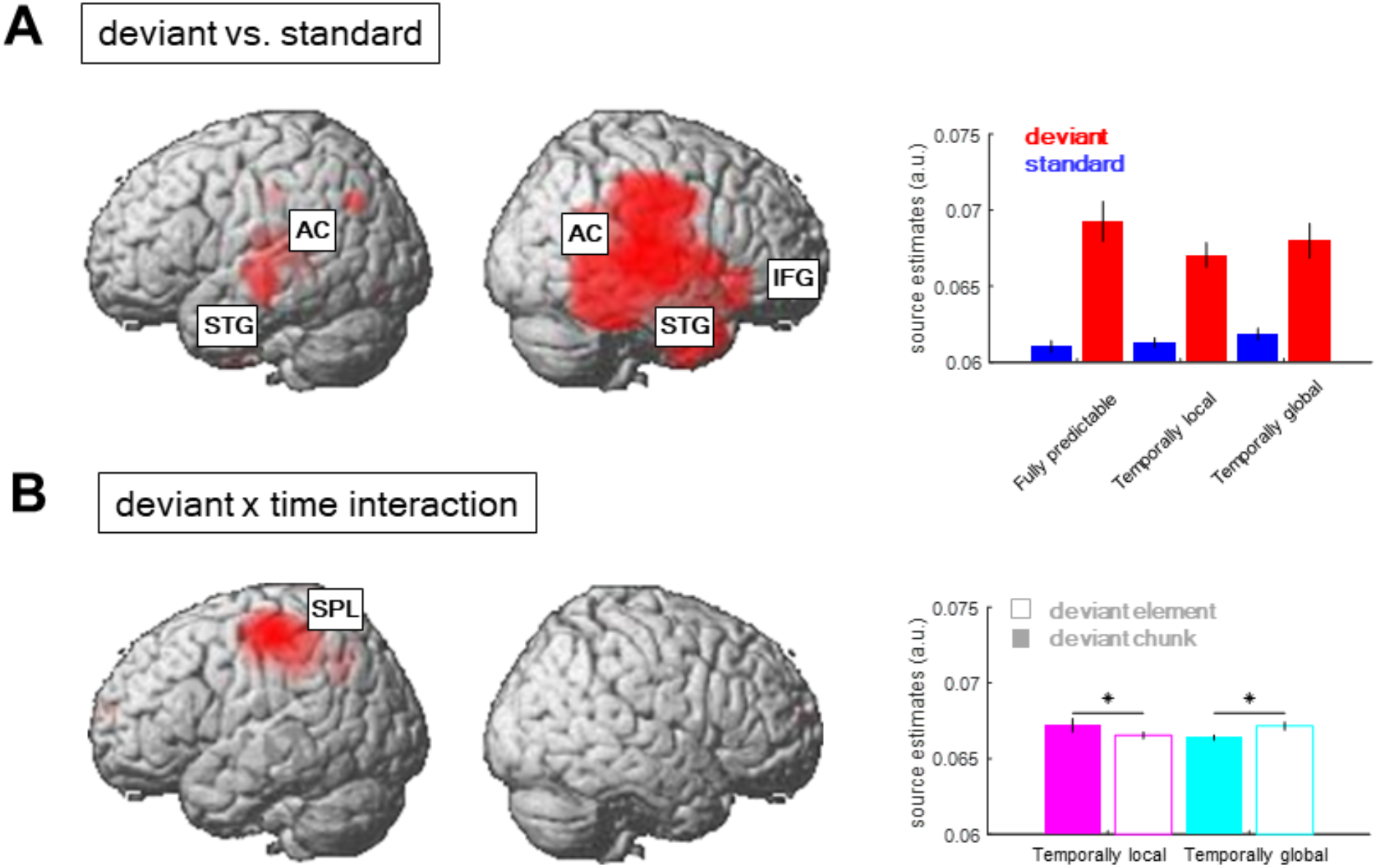
Source reconstruction. **(A)** Regions showing a significant main effect of content-based predictions (deviant vs. standard). Inset shows average source estimates per condition. Error bars denote SEM across participants. **(B)** Regions showing a significant contextual interaction effect between content-based predictions (deviant element vs. deviant chunk) and temporal predictions (temporally local vs. temporally global). Figure legend as in (A).

The difference between source estimates associated with deviants and standards was localized to a large network of regions (see Table 1 for full results), including bilateral auditory cortex (AC) and superior temporal gyri (STG) and the right inferior frontal gyrus (IFG). On the other hand, the interaction effect between deviant types and temporal conditions was localized to a spatially confined cluster in the left superior parietal lobule (SPL; see Table 1). A post-hoc analysis revealed that, in this cluster, element deviant responses presented in the temporally-local condition were associated with weaker source estimates than chunk deviant responses presented in the temporally-local condition (T_max_ = 3.67, Z_max_ = 3.46, p_FWE_ = 0.009, small-volume corrected). Similarly, chunk deviant responses presented in the temporally-global condition were associated with weaker source estimates than element deviant responses presented in the temporally-global condition (T_max_ = 5.79, Z_max_ = 5.11, p_FWE_ = 0.003, small-volume corrected). Thus, while the deviant processing could be linked to a wide network of auditory and frontal regions, deviants presented in the corresponding temporal predictability conditions (e.g., element deviants in the temporally-local context) were associated with a relative decrease of left parietal activity.

**Table 1.** Source reconstruction results. Summary of significant clusters showing differences between conditions.

| Effect | Cluster label | Peak MNI coords | $F_{\max}$ | $Z_{\max}$ | Voxel extent | $p_{\text{FWE}}$ |
| --- | --- | --- | --- | --- | --- | --- |
| Deviant vs. standard | Right transverse temporal gyrus / auditory cortex (AC) | 48 -20<br>12 | 53.9<br>9 | 4.8<br>3 | 2050<br>8 | < 0.001 |
|  | Right superior temporal gyrus (STG) | 44 -48<br>12 | 40.1<br>5 | 4.4<br>2 |  |  |
|  | Right inferior frontal gyrus (IFG) | 40 26<br>-6 | 34.5<br>2 | 4.2<br>0 |  |  |
|  | Left transverse temporal gyrus / auditory cortex (AC) | -38 -28<br>12 | 34.3<br>1 | 4.2<br>0 | 2177 | 0.003 |
|  | Left superior temporal gyrus (STG) | -60 -20<br>-8 | 31.1<br>9 | 4.0<br>6 |  |  |
| ("element" vs. "chunk" deviant) x (temporally-local vs. temporally-global) | Left superior parietal lobule (ISPL) | -26 -40<br>46 | 49.3<br>7 | 5.8<br>2 | 3073 | 0.003 |

### Dynamic causal modeling

To infer the most likely effective connectivity patterns underlying the observed ERP results, we used the six main cortical regions identified in the source reconstruction results as regions of interest (ROIs) to build a generative model of the ERP data. A fully interconnected model, fitted to each participants’ ERP data, explained on average 71.03% of the ERP variance (SEM across participants 2.81%).

Bayesian model reduction was used to obtain connectivity and gain parameters of a range of reduced models, in which only a subset of parameters were allowed to be modulated by the two conditions (deviant vs. standard; interaction deviant element/chunk x temporally-local/global). Using this procedure, we identified a single winning model, in which “what” predictions (deviant vs. standard) modulated all types of connections (ascending, descending, lateral, and intrinsic), while the interaction between “what” and “when” predictions modulated three out of four types of connections (ascending, lateral, and intrinsic). The difference between the free-energy approximation to log-model evidence between the winning model and the next-best model (i.e., log Bayes factor) was 5.6615, corresponding to very strong evidence for the winning model (>99% probability). Therefore, the resulting Bayesian model average, implemented to integrate model parameter estimates from the entire model space, was mostly informed by the single winning model.

The posterior parameter estimates of the Bayesian model average are plotted in Fig. 5A and reported in Table 2. The results revealed that deviant processing (as opposed to standard processing) significantly increased nearly all connectivity estimates (probability of increase >99.9% for all parameters), corresponding to an increase in excitatory ascending connectivity and in inhibitory descending and intrinsic (gain) connectivity - with the exception of the intrinsic self-inhibition in the left SPL region, which was significantly decreased following deviant processing, as well as the bidirectional connectivity between the left SPL and right IFG, which was not affected by deviant processing.

**Table 2.**
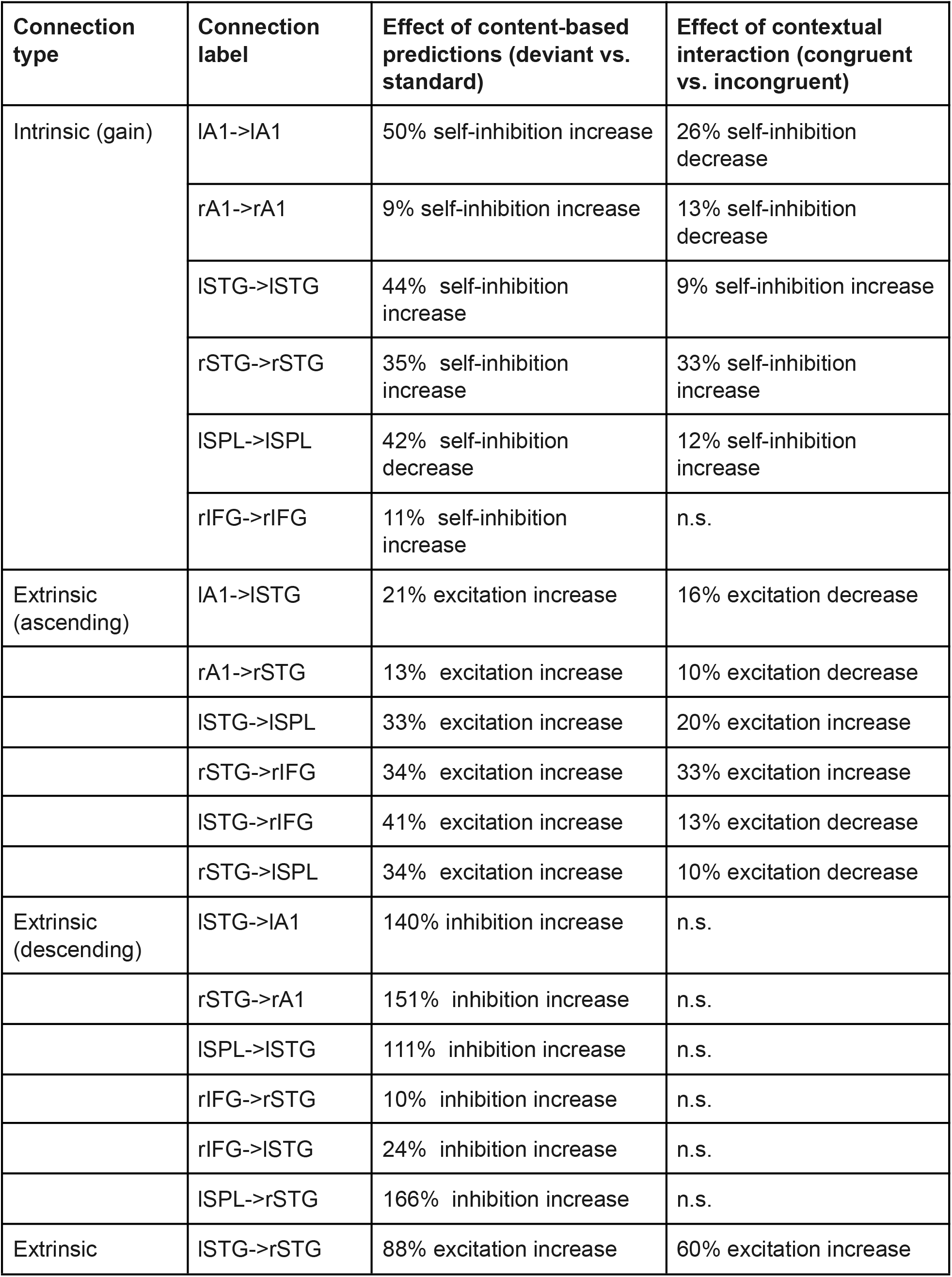
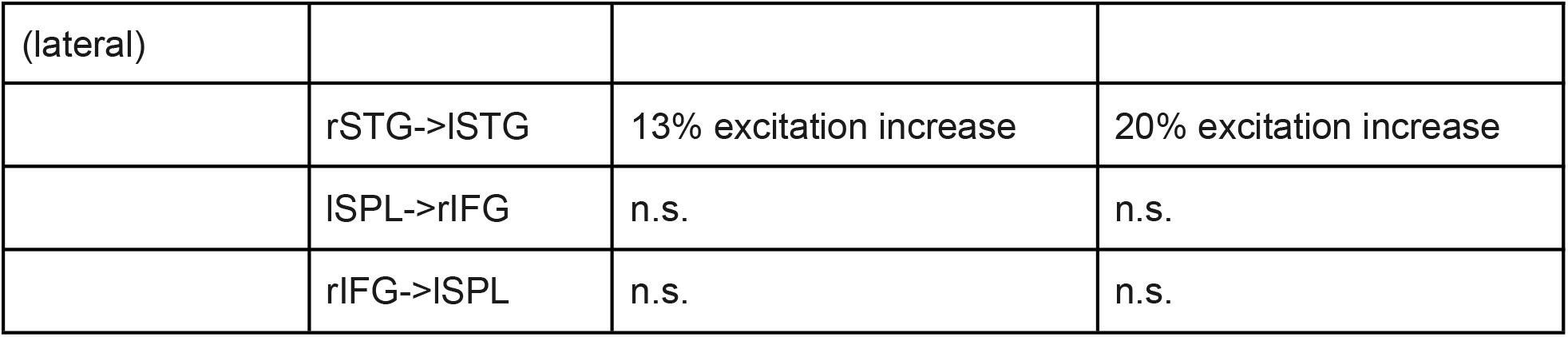
Dynamic causal modeling results. Summary of significant condition-specific effects on connectivity estimates.

| **Connection type** | **Connection label** | **Effect of content-based predictions (deviant vs. standard)** | **Effect of contextual interaction (congruent vs. incongruent)** |
| --- | --- | --- | --- |
| Intrinsic (gain) | IA1->IA1 | 50% self-inhibition increase | 26% self-inhibition decrease |
|  | rA1->rA1 | 9% self-inhibition increase | 13% self-inhibition decrease |
|  | ISTG->ISTG | 44% self-inhibition increase | 9% self-inhibition increase |
|  | rSTG->rSTG | 35% self-inhibition increase | 33% self-inhibition increase |
|  | ISPL->ISPL | 42% self-inhibition decrease | 12% self-inhibition increase |
|  | rIFG->rIFG | 11% self-inhibition increase | n.s. |
| Extrinsic (ascending) | IA1->ISTG | 21% excitation increase | 16% excitation decrease |
|  | rA1->rSTG | 13% excitation increase | 10% excitation decrease |
|  | ISTG->ISPL | 33% excitation increase | 20% excitation increase |
|  | rSTG->rIFG | 34% excitation increase | 33% excitation increase |
|  | ISTG->rIFG | 41% excitation increase | 13% excitation decrease |
|  | rSTG->ISPL | 34% excitation increase | 10% excitation decrease |
| Extrinsic (descending) | ISTG->IA1 | 140% inhibition increase | n.s. |
|  | rSTG->rA1 | 151% inhibition increase | n.s. |
|  | ISPL->ISTG | 111% inhibition increase | n.s. |
|  | rIFG->rSTG | 10% inhibition increase | n.s. |
|  | rIFG->ISTG | 24% inhibition increase | n.s. |
|  | ISPL->rSTG | 166% inhibition increase | n.s. |
| Extrinsic | ISTG->rSTG | 88% excitation increase | 60% excitation increase |
| (lateral) |  |  |  |
|  | rSTG->ISTG | 13% excitation increase | 20% excitation increase |
|  | ISPL->rIFG | n.s. | n.s. |
|  | rIFG->ISPL | n.s. | n.s. |

**Figure 5.**
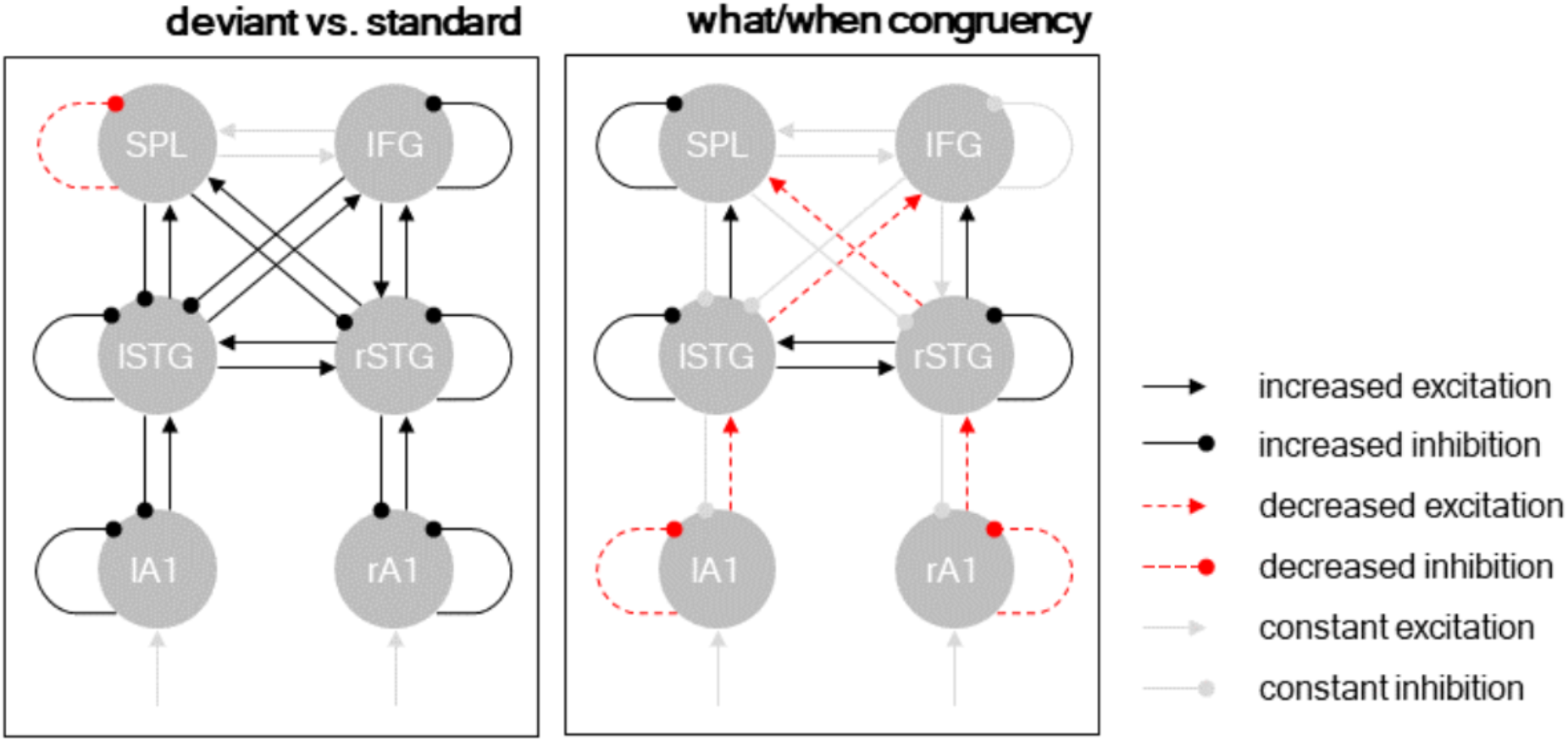
Dynamic causal modeling. Posterior model parameters. Separate panels show different condition-specific effects. Black arrows: excitatory connections; red arrows: inhibitory connections; solid lines: condition-specific increase; dashed lines: condition-specific decrease. Significant parameters (p < 0.001) shown in black/red, remaining connections (constant excitation/inhibition) shown in gray.

The interaction between deviant type (deviant element vs. deviant chunk) and temporal predictability (temporally-global vs. temporally-local) modulated a more nuanced connectivity pattern. At the hierarchically lower level (between A1 and STG), deviants processed in a temporally congruent condition (i.e., deviant elements in the temporally-local condition, and deviant chunks in the temporally-global condition) decreased excitatory ascending connectivity from A1 to STG and inhibitory self-connectivity in A1. Conversely, at the hierarchically higher level (between STG and the fronto-parietal regions), deviants processed in a temporally congruent condition increased excitatory ascending connectivity from STG to SPL/IFC and inhibitory self-connectivity in the STG. Furthermore, deviants processed in a temporally congruent condition (1) increased lateral connectivity between the left and right STG, (2) decreased cross-hemispheric ascending connectivity between the STG regions and the fronto-parietal regions, and (3) increased self-inhibition in the left SPL region.

## Discussion

In the present study, we found that “when” predictions modulate MMR to violations of “what” predictions in a contextually specific fashion, such that more local “when” predictions modulated responses to single deviant elements, while more global “when” predictions modulated responses to deviant chunks, indicating a congruence effect in the processing of “what” and “when” predictions at different contextual levels in the auditory system. The authors interpret this as levels of processing hierarchy, however it is worth noting that alternate interpretations of this effect could rely on simple contextual pairings such as position effects within the sequence. While “what” and “when” kinds of predictions showed interactive effects for both levels, both interaction effects (e.g. chunk-rate/deviant chunk and element-rate/deviant element) were associated with similar spatiotemporal patterns of EEG evoked activity modulations, and linked in the DCM analysis to a widespread connectivity increase at relatively late stages of cortical processing (between the STG and the fronto-parietal network). These findings suggest that the integration of “what” and “when” predictions, while contextually specific, is mediated by a shared and distributed cortical network.

Deviant responses to “what” prediction violations within melodic sequences and tone contours are well documented, having been used to explore a variety of phenomena in the auditory system (see Yu et al., 2015 for a partial review). Deviant tones within familiar musical scales have been found to elicit higher MMR amplitudes compared to those of unfamiliar scales are tones presented without a scale structure (Brattico et al., 2001), as well as higher deviant responses to out-of-scale notes in unfamiliar melodies (Brattico et al., 2006). Deviant responses to manipulated musical characteristics within melodic sequences (e.g. timing, pitch, transposition, melodic contour) have similarly been demonstrated in musician and non-musician groups (Tervaniemi et al., 2014, 2014; Vuust et al., 2011). In predictive coding frameworks, such evoked responses can be understood in the context of prediction error, wherein bottom-up error signaling triggers the adjustment of higher-level models of the stimulus train formed as a result of perceptual learning during repeated stimulus presentation (Garrido et al., 2009). Such hierarchical relationships have been quantified using DCM (Auksztulewicz and Friston, 2016), and are consistent with our analysis of the evoked responses observed herein. The resultant model shows increased connectivity throughout the network, consistent with increased error signaling (ascending connections), predictive template updates (descending connections), and gain connectivity evident in a decrease in gain following predictions errors. Our source reconstruction was equally consistent with existing literature revealing bilateral activity in the primary auditory cortex (A1) and higher-order auditory regions in the superior temporal gyrus (STG), as well as the right inferior frontal gyrus (IFG) (Garrido et al., 2008; Giroud et al., 2020).

Turning to “when” predictions, the results of our frequency domain analysis show that the EEG spectrum largely follows that of the stimulus spectrum. However, ITPC peaks at the pair-tone rate of ‘fully predictable’ and ‘temporally local’ sequences are significantly larger than neighboring frequencies, which is not the case in the stimulus spectrum, indicating that ITPC peaks do not just follow the stimulus spectrum but also reflect the neural processing of sequence structures at higher levels (e.g. chunking (Kotz et al., 2018)). Indeed, our behavioral results show faster reaction times in temporally predictable conditions, supporting the notion of neural entrainment to stimulus periodicity, results which mirror previous behavioral studies (Morillon et al., 2016). We found that the EEG-based ITPC response at tone-rate is stronger near central electrodes, with results consistent with existing EEG studies (Ding et al., 2017). Additionally, the chunk-rate effect is predominantly present in the right hemisphere, suggesting that the contextual structure of non-linguistic sequences can be entrained by parallel neural activity in different regions at distinct time scales - consistent with existing research (Giroud et al., 2020). Interestingly, the ITPC differences between conditions (temporally-local vs. global) emerged during the experiment in chunk-rate peaks, but not in telement-rate peaks, suggesting that rapid learning could modulate neural entrainment to auditory sequences with different regularities at the chunk-rate level. Similarly, a previous study (Moser et al., 2021) found significant differences in non-linguistic triplet-rate ITPC peaks between structured and random conditions, occurring during early exposure. This ITPC difference suggests a fine shift in sequence encoding, with different regularities from single elements to integrated chunks. Notably, we also found correlations between the ITPC difference conditions and the congruence effect of ERP amplitude, indicating a mutual network between neural entrainment and prediction.

In addition to their dissociable main effects on neural activity, “what” and “when” predictions modulated element-evoked response amplitude interactively and in a contextually specific manner, such that faster “when” predictions amplified MMRs to less complex “what” prediction violations (single elements), while slower “when” predictions amplified MMRs to more complex “when” prediction violations (chunks). These findings extend the result of previous studies, which showed that “when” predictions modulate MMR amplitude (Jalewa et al., 2021; Lumaca et al., 2019; Takegata and Morotomi, 1999; Todd et al., 2018; Yabe et al., 1997), by showing that these modulatory effects are specific with respect to the complexity of “what” predictions. Dynamic causal modeling of our ERP data showed partially dissociable connectivity patterns between the main effect of “what” predictions (i.e., all deviants vs. all standards), which increased recurrent connectivity throughout the network (Auksztulewicz and Friston, 2015; Fitzgerald et al., 2021; Garrido et al., 2008), and “what”/”when” interactive effects, which had a more nuanced pattern of effects on neural activity. Specifically, congruent “what” and “when” predictions decreased recurrent connectivity at lower parts of the network (between A1 and the STG), while at the same time increasing recurrent connectivity at higher parts of the network (between STG and the fronto-parietal regions). Previous DCM work has shown similar dissociations between processing deviants based on violations of relatively simple predictions vs. complex contextual information, indicating the higher-order regions as sensitive to complex prediction violations (Fitzgerald et al., 2021). Additionally, in the current results, the main effect of “what” predictions and the contextually specific integration of “what” and “when” predictions had opposing effects on the neural gain estimates for the left SPL region, which displayed decreased self-inhibition (increased gain) following deviant processing but increased self-inhibition (decreased gain) following prediction integration. These results mirror our source reconstruction, wherein deviants presented in congruent temporal conditions were associated with decreased left parietal activity, and imply the left parietal cortex - recently shown to mediate the integration of “what” and “when” information in speech processing (Orpella et al., 2020) - in the more general process of integrating “what” and “when” predictions also for non-speech stimuli. It is worth noting that while “when” predictions did not elicit a significant main effect on ERP amplitude, it is possible this finding may have resulted from design constraints, as all conditions contained only “what” (repetition detection) tasks, suggestive of previous studies on the role of attention in parallel temporal and mnemonic predictive processing (Lakatos, et al., 2013, Wollman and Morillon, 2018).

Previous studies have shown that the processing of musical information requires predictive mechanisms for timing of content of auditory events, and that such predictions can have modulatory effects at different cortical levels when presented within the framework of melodic expectation (Di Liberto et al., 2020; Royal et al., 2016). Musical stimuli presents us with an intriguing opportunity to investigate predictive coding mechanisms, as the statistical regularities within musical frameworks are well defined and intrinsically learned. In particular, such structures allow us to disassociate “what” and “when” predictions while keeping other elements of a stimulus stream intact across manipulations and trials. Studies have demonstrated an early right anterior negativity (ERAN) in contexts where musical syntax has been violated, as opposed to the comparatively low-level acoustic diavations that elicit a MMN response (see Koelsch et al., 2019 for review). Because the presence of musical syntax violations require knowledge acquired through long-term repeated exposure to music, long-term memory recall is involved in establishing those regularities. The role of memory in syntactical prediction violation is an avenue ripe for further investigation, and future studies may wish to extend our paradigm to further probe the observed late-series ITPC pair-rate differences in that context. Furthermore, since “what” and “when” predictions are also ubiquitous in other stimulus domains - most prominently in speech perception - future research should test whether similar contextual specificity of “what” and “when” predictions as observed here also governs speech processing.

## Acknowledgments

This work has been supported by the European Commission’s Marie Skłodowska-Curie Global Fellowship (750459 to R.A.) and a grant from the European Commission / Hong Kong Research Grants Council Joint Research Scheme (9051402 to R.A. and J.S.).

## Notes

### Competing Interest Statement

The authors have declared no competing interest.

